# Genome-wide mapping of G-quadruplex structures with CUT&Tag

**DOI:** 10.1101/2021.04.25.441312

**Authors:** Jing Lyu, Rui Shao, Simon J Elsässer

## Abstract

Single-stranded genomic DNA can fold into G-quadruplex (G4) structures or form DNA:RNA hybrids (R loops). Recent evidence suggests that such non-canonical DNA structures affect gene expression, DNA methylation, replication fork progression and genome stability. When and how G4 structures form and are resolved remains unclear. Here we report the use of Cleavage Under Targets and Tagmentation (CUT&Tag) for mapping native G4 in mammalian cell lines at high resolution and low background. Mild native conditions used for the procedure retain more G4 structures and provide a higher signal-to-noise ratio than ChIP-based methods. We determine the G4 landscape of mouse embryonic stem cells (mESC), observing widespread G4 formation at active promoters, active and poised enhancers. We discover that the presence of G4 motifs and G4 structures distinguishes active and primed enhancers in mESCs. Further, performing R-loop CUT&Tag, we demonstrate the genome-wide co-occurence of single-stranded DNA, G4s and R loops, suggesting an intricate relationship between transcription and non-canonical DNA structures.

## INTRODUCTION

G-quadruplex (G4) structures are composed of three or more stacked G-quartets. Four guanine bases can form a planar G-quartet via Hoogsten hydrogen bonds, and the stacking of G-quartets is stabilized by monovalent cations, preferably K^+^. The DNA backbones of the guanines run parallel or antiparallel along the stack, connected by short, sometimes also longer, single-stranded DNA loops (1). Moreover, G-quadruplex DNA can form intramolecularly or intermolecularly: Four GGG-repeats connected by short linkers on the same DNA molecule form the canonical G4 motif, but G4s have also been shown to fold with as few as two or more than three guanine quartets or non-G bases breaking up the G-repeat. Further, GGG-repeats distributed on both strands of a DNA duplex can form inter-strand G4s, and it has been proposed that G4s can also form in trans between two DNA molecules.

First demonstrated in prokaryotes, G4 structures occurring at promoter regions are implicated in gene regulation (2). G4 DNA further was found in telomere regions in Stylonichia lemnae (3, 4). The human genome contains up to half a million predicted G-quadruplex forming sequences (PQS), most of which are found in promoter regions/CpG islands, G-rich tandem repeat regions and telomeres. Experimental evidence suggests that G4 structures are also enriched in telomeric and sub-telomeric repetitive DNA, ribosome DNA, promoter regions, tandem repeats in mammalian cells (5, 6). RNA can also form G4 structures. G4-forming sequences are significant underrepresentation in the coding strand of exonic regions, which indicates that RNA G4 structures have been selected against in evolution (7, 8). Nevertheless, RNA G4 structures are thought to regulate mRNA metabolism, and RNA may form hybrid G4s with DNA (9, 10).

G4 motif is enriched in the promoter region of oncogenes, such as *c-MYC, KRAS* and *c-kit* (11-13), raising the possibility that G4 structures participate in disease mechanisms or could be exploited as drug targets (14-16). Owing to the unique stacking of the G quartets, many specific small molecule ligands have been developed. Intercalating between the G-quartets, G4 ligands have found applications in detecting G4 structures, stabilizing or inducing them (14-20). TMPyP4 has been shown to repress *c-MYC* and the expression of telomerase reverse transcriptase in mice (21), and consistent findings were reported in leukemic cells (22). Pyridostatin (PDS) was shown to induce telomere dysfunction and inhibit cancer growth (23). One study suggested that PDS predominantly binds non-telomeric regions at lower concentration (24). Many G4 ligands also intercalate into duplex DNA and/or induce other non-canonical DNA structures, thus raising the question if observed phenotypes, in particular cell toxicity, could be unequivocally linked to G4s (25). Recently, a new G4 ligand PDC12 was screened, and it was demonstrated that PDC12 was less toxic and more specific than PDS due to its low-molecular-weight (25). Cancer cells have been shown to be relying on elevated rDNA transcription. Several inhibitors for rRNA synthesis have been developed in recent years, such as, CX-3543 (26-29), CX-5461 (30-32), and BMH-21 (33-35). These small molecules have been shown to bind rDNA G-quadruplex and disrupt nucleolin/rDNA G-quadruplex complexes, thereby inhibiting RNA polymerase I elongation and rRNA synthesis, and inducing apoptosis in cancer cells. Therefore, CX-3543, CX-5461 and BMH-21 exhibit potent antitumor activity in cancer models. It has also been shown that endogenous G4 levels can predict sensitivity of tumors to G4 ligands (36).

While these studies show the potential of G4 stabilizing agents to disrupt various DNA-dependent processes, the role of G4 in endogenous control of transcription and replication remains unclear. Elegant studies in chicken cells show that individual G4 structures can represent roadblocks to replication (25, 37, 38), yet such effect has not been confirmed on a genome wide level.

Methods to precisely map DNA G-quadruplexes in the genome are crucial to study the functions and regulations of G4 in physiological and pathological processes. Genome-wide chromatin-immunoprecipitation and sequencing (ChIP-seq) based mapping methods have been described, using either a G4-specific antibody in an optimized ChIP-Seq protocol (36, 39-41) or an artificial 6.7 kDa G4 probe (G4P) protein for G4 binding and capture (42).

G4 ChIP-seq is a method which couples formaldehyde-fixed G4 DNA immunoprecipitation using a G4-specific antibody (most commonly the phage-display derived monoclonal BG4 antibody (43)) with high-throughput sequencing (39). Using G4 ChIP-Seq, approximately 10,000 DNA G4 structures were mapped in the human genome, enriched in promoters and other regulatory, nucleosome free regions, and G4 formation was correlated with elevated transcriptional activity (39, 41). A further study suggests that G4 formation may contribute to the protection of CpG islands from cytosine methylation, by sequestering DNMT1 (40). These results highlight the utility of genome-wide G4 ChIP-Seq to determine genome-wide distribution of G4s. However, formaldehyde fixation used in G4 ChIP-Seq to maintain the structures in the presence of various detergents used during the ChIP procedure may not protect all G4 structures and additionally bears the risk of masking G4 recognition by the antibody, thus reducing specific signal. Moreover, shearing of the crosslinked chromatin by sonication may dissolve or distort G4 structure and also affect antibody recognition. Thus, alternative approaches have been sought to capture G4s under more native conditions: the G4P-ChIP method takes advantage of a peptide from the DHX36 helicase with high affinity and specificity for G4 structures, whereas D1 ChIP used a mononclonal antibody-GFP fusion protein (42, 44). However, both methods rely on expression of the G4 binder in cells, thus requiring the generation of stable cell lines, and it cannot be excluded that the expression of G4 binding proteins affects G4 biology, e.g. by competing with endogenous binding proteins or helicases.

Recently, cleavage under targets and tagmentation (CUT&Tag) has been reported to map chromatin features in permeabilized nuclei using antibody-targeted Tn5 tagmentation (45), In CUT&Tag, cells or nuclei are permeabilized under mild native conditions. Here, we combined G4 antibody-based detection of genomic G4s with CUT&Tag technology and established a native G4 mapping method called G4 CUT&Tag. Compared to other G4 mapping methods, G4 CUT&Tag provide superior signal-to-noise and reliability to detect bona-fide G4s. We applied G4 CUT&Tag to mouse embryonic stem cells, finding widespread G4 formation at active and enhancers. Together, our study provides a simple and fast methodology that accurately maps DNA G4s across the genome.

## RESULTS

### Systematic comparison of G4 CUT&Tag with other G4 mapping methods

We established G4 CUT&Tag starting with version 2 of CUT&Tag (45) with minor modifications (see Materials and Methods). To systematically compare the G4 CUT&Tag with other G4 mapping methods, we performed G4 CUT&Tag in cell lines for which prior genome-wide datasets had been generated. In parallel, we reprocessed the raw data for G4 ChIP-seq (41), G4P-ChIP (42) through the same bioinformatic pipeline as our G4 CUT&Tag data. G4 CUT&Tag demonstrated improved raw data quality compared to both existing methods, yielding tenfold higher fraction of reads under peaks (FriP) than G4 ChIP-seq and by sixfold higher FriP than G4P-ChIP (Figure 1A). Fingerprint plots also demonstrated the higher signal-to-noise ratio of G4 CUT&Tag (Figure 1B, C). Examples show the large pileup of G4 CUT&Tag on promoters with predicted G4 sequences (Figure 1D). We further evaluate the overlap of peaks identified by G4 CUT&Tag with peaks identified by G4 ChIP-seq or G4P-ChIP. To allow an unbiased comparison, the top 10,000, top 5000, top 1000 highest-scoring peaks identified by each method were extracted. Comparing G4 CUT&Tag and G4 ChIP-seq, 33% of top 10,000 peaks, 38% of top 5000 peaks and 30% of top 1000 peaks were identified by both methods, respectively (Figure 1E; Supplementary Figure 1). Of top peaks identified by G4 CUT&Tag and G4P-ChIP, 45.83% of top 10,000 peaks, 32.76% of top 5000 peaks and 11.7% of top 1000 peaks are identified by both methods, respectively (Figure 1F; Supplementary Figure 1A, B). While the relatively small overlap would suggest a substantially different distribution of peaks, assessing the enrichment amongst shared and unique peaks showed that most called peaks, no matter from which method, showed enrichment by the methods under comparison (Figure 1E, F; Supplementary Figure 1C, D). Nevertheless, differences existed between the quality of the peaks called: Of top peaks that were unique to each method, the G4 CUT&Tag protocol generated a higher percentage of peaks that match predicted G4 sequences (PQS, including inter-strand and intra-strand G4s) (Figure 1G, H). This further demonstrates that G4 CUT&Tag protocol allows mapping G4 structures with higher confidence than alternative methods. Our G4 CUT&Tag data was generated from 1×10^5^ cells. In contrast, cell number used in the previous G4 mapping methods is 1×10^7^ in G4 ChIP-seq and 1×10^8^ in G4P-ChIP. Thus, in summary, G4 CUT&Tag enabled a dramatic increase in the signal-to-noise ratio at much reduced input requirements, as compared to G4 ChIP-seq or G4P-ChIP.

**Figure 1:**
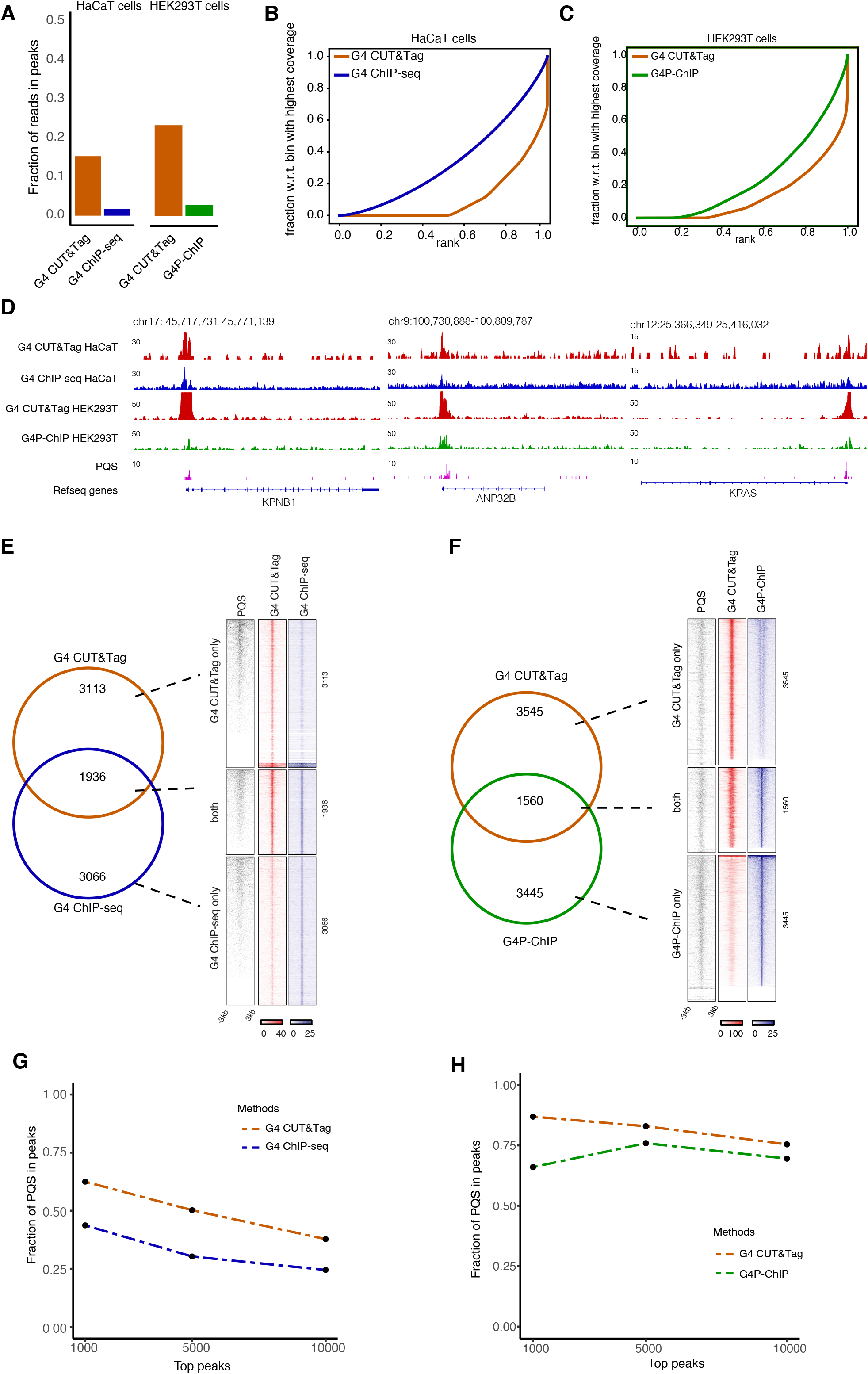
Comparison of G4 CUT&Tag to other G4 mapping methods, G4 ChIP-seq and G4P-ChIP. (**A**) Fraction of reads in G4 peaks comparison between G4 CUT&Tag library generated in this study and from public G4 ChIP-seq dataset in HaCaT cells, or public G4P-ChIP dataset in HEK293T cells (41, 42). (**B**) Fingerprint (cumulative read count sum by ranked bins) plot of G4 CUT&Tag and G4 ChIP-seq in HaCaT cells (41). (**C**) Fingerprint (cumulative read count sum by ranked bins) plot of G4 CUT&Tag and G4P-ChIP in HEK293T cells (42). (**D**) Genome browser view of G4 signals at example loci. The RPGC-normalized (1x Genome Coverage) tracks of G4 CUT&Tag and G4 ChIP-seq in HaCaT cells, or G4 CUT&Tag and G4P-ChIP in HEK293T cells are shown on the same y-axis scale. (**E)** Overlap of top 5000 G4 CUT&Tag and G4 ChIP-seq peaks. Total PQS, G4 CUT&Tag and G4 ChIP-seq density heatmaps for peaks classified as G4 CUT&Tag only (n = 3113), both G4 CUT&Tag and G4 ChIP-seq (n = 1936) or G4 ChIP-seq only (n = 3066). (**F**) Overlap of top 5000 G4 CUT&Tag and G4P-ChIP peaks. Total PQS, G4 CUT&Tag and G4P-ChIP density heatmaps for peaks classified as G4 CUT&Tag only (n = 3545), both G4 CUT&Tag and G4P-ChIP (n = 1560) or G4P-ChIP only (n = 3445). (**G**) Comparison of fraction of PQS (canonical or non-canonical) in top 10,000, top 5000, and top 1000 G4 peaks between G4 CUT&Tag and G4 ChIP-seq in HaCaT cells. (**H**) Comparison of fraction of PQS (canonical or non-canonical) in top 10,000, top 5000, and top 1000 G4 peaks between G4 CUT&Tag and G4P-ChIP in HEK293T cells.

### Characterization of G4s in mouse embryonic stem cells

Next, we performed G4 CUT&Tag in mouse embryonic stem (mES) cells. mES cells are primary pluripotent stem cells featuring an open and dynamic chromatin structure that maintains an uncommitted state poised for differentiation (46). G4 structures have been proposed to play a regulatory role in animal development (47). The G4 landscape of mES cells has not been elucidated to date.

mES G4 CUT&Tag was robust and reproducible across triplicate experiments (Figure 2A, Supplementary Figure 2A,B). 9186 high-confidence G4 CUT&Tag peaks were identified by intersecting peaks called from the three replicates, showing good overlap with canonical and non-canonical (trans-strand) PQS (Figure 2B). To further validate the specificity of G4 CUT&Tag for G4 motifs, we interrogate the top 1000, top 500 and top 200 G4 peaks for recurring motifs using MEME suite and find that G-rich sequences are highly prevalent among the top peaks (Figure 2C). G4s predominantly associated with active promoters (61%) and enhancers (13%), and coincided with open chromatin, H3K4me3, H3K27ac and H3K4me1 (Figure 2B, D). Importantly, a CUT&Tag control experiment with a H3K27me3 antibody showed enrichment at bivalent but not active TSS, demonstrating that the strong enrichment over accessible sites is not due to an artificial bias of the methodology. A small proportion (3%) of G4s also coincided with repressed or bivalent promoters, enriched for H3K27me3 (Figure 2B, C, Supplementary Figure 2C, D). G4 profiles on gene-coding regions showed a strong peak around the transcription start sites (TSS), while largely absent in the gene body despite the presence of PQS (Supplementary Figure 2B). Efficient chromatin maintenance during RNA Polymerase passage, relying on Spt6, FACT and other histone chaperones is known to maintain dense nucleosome occupancy in gene bodies (48), thus likely disfavoring formation of G4. Promoters of non-expressed genes, exhibiting neither H3K4me3 or H3K27me3 showed the lowest G4 signal (Supplementary Figure 2D). As known in human cells (40), G4s correlated with CpG density, and anticorrelated with CpG methylation (Supplementary Figure 2E-H).

**Figure 2:**
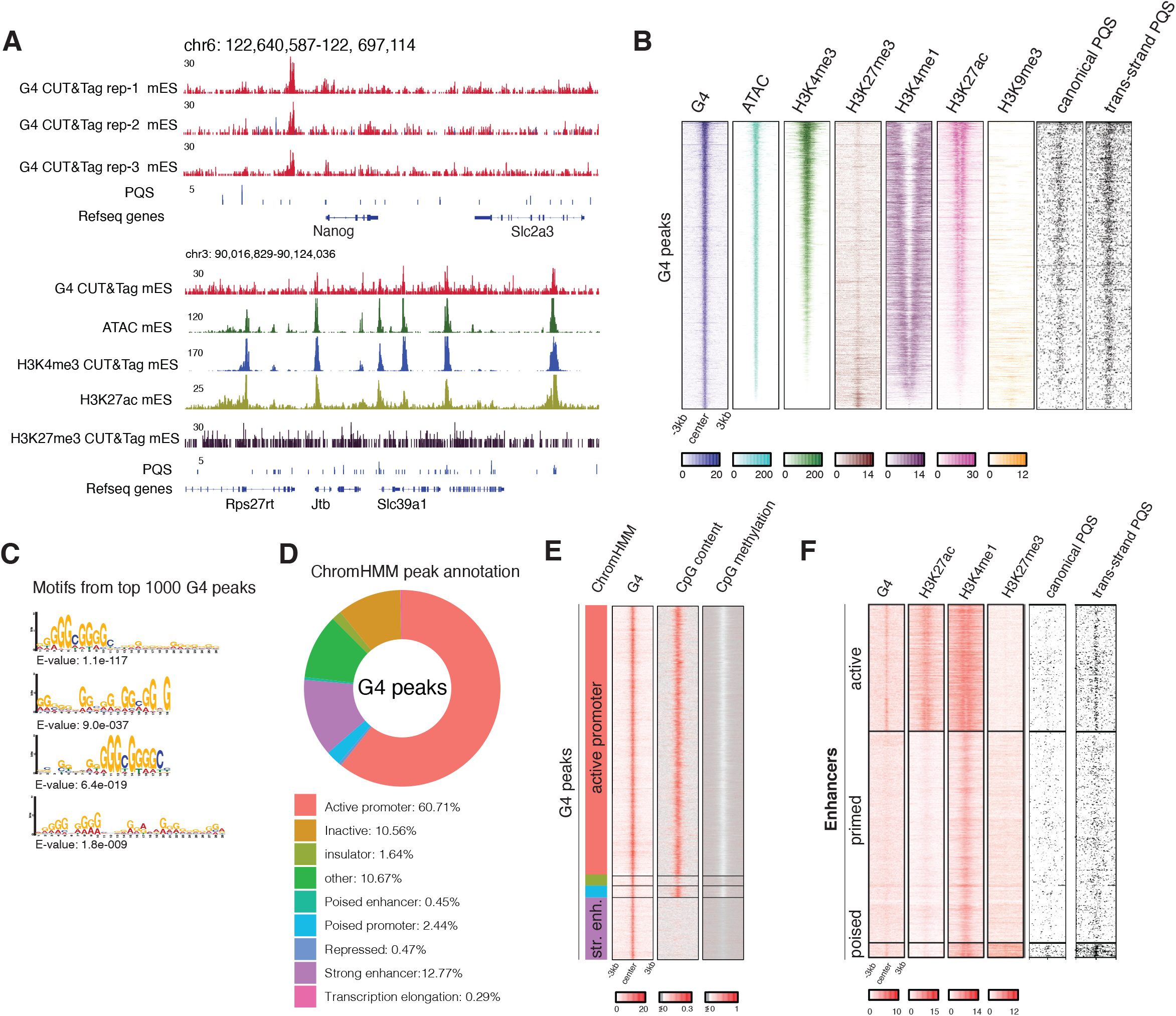
G4 landscape in mouse embryonic stem cells. (**A**) Genome browser view showing triplicate G4 CUT&Tag experiment in mES cells at *Nanag* locus, and comparison of G4 CUT&Tag, H3K4me3 and H3K27me3 CUT&Tag from the same cells, as well as published ATAC-seq (63) and H3K27ac ChIP-Seq (64). (**B**) Density heatmaps of the same tracks at high-confidence (shared between three replicates) G4 peaks in mES cells. Prediction of canonical and non-canonical (trans-strand) PQS are shown. (**C**) Motif discovery using MEME suite from top 1000 G4 peaks shows typical canonical and non-canonical G4 patterns. (**D**) Annotation of high-confidence G4 peaks with different functional genomic features as defined by ChromHMM (65). (**E**) Density heatmaps of G4 CUT&Tag, CpG content and methylation and G4 peaks grouped by ChromHMM annotation as in (D). (**F**) Density heatmaps of G4 CUT&Tag, H3K27ac, H3K4me1, H3K27me3, canonical and non-canonical PQS at active enhancer (active), primed enhancer (primed) and poised enhancer (poised) regions (49).

Enhancers are much less GC-rich than promoters, however they were still amongst the most abundant G4-enriched regions (Figure 2E). To further investigate this, we assessed G4s across a reference list of active, primed or poised enhancers (49): G4s were found at active and poised, but not primed enhancers (Figure 2F). Notably, enhancers lacked canonical G4-motifs. Instead, non-canonical G4 motifs were underlying G4s at both active and poised enhancers (Figure 2F), and G4 motifs were essentially absent from primed enhancers. Thus, surprisingly, the presence/absence of G4 motifs is a differentiating feature of active and primed enhancers (Figure 2F).

### G4 CUT&Tag signals are sensitive to single-strand specific nuclease digestion

G4s formed by one DNA strand leave the C-rich opposite strand single-stranded (Figure 3A). Indeed, methods to map single-stranded DNA (ssDNA) reported widespread ssDNA formation at active TSS (50, 51). A recent study using kethoxal-assisted single-stranded DNA sequencing (KAS-seq) showed that ssDNA regions were maintained even after inhibition of transcriptional elongation or depletion of RNA Polymerase II, suggesting that the ssDNA regions were maintained by a mechanism independent of continuous transcription (50). Indeed, KAS-seq indicated ssDNA at G4 peaks (Figure 3D), and we found a positive correlation between KAS-seq and G4s across TSS (Supplementary Figure 3A). We wondered if we can corroborate the co-existence of G4 and ssDNA, and thus indirectly validate G4 CUT&Tag signals, by applying a single-strand specific endonuclease to permeabilized cells, before subjecting to G4 CUT&Tag. Mung Bean nuclease is a single-strand DNA/RNA-specific endonuclease (52). Digestion of ssDNA by Mung Bean Nuclease should release G4 structures and/or inhibit PCR amplification of the G4-associated, tagmented DNA (Figure 3A). mES cells were pretreated with 100 units and 150 units of Mung Bean nuclease respectively. G4 CUT&Tag signals decreased at TSS regions after Mung Bean nuclease treatment, and analysis of high-confidence G4 peaks confirmed that Mung Bean treatment led to substantial reduction of G4 signals across all G4 (Figure 3B-D). After 100 units and 150 units of Mung Bean nuclease treatment, G4 CUT&Tag peak numbers decreased by 78.6% and 88.3% respectively (Figure 3C). Thus, digestion of ssDNA removes G4-specific reads from the CUT&Tag, leaving unspecific background signal largely unaffected (Figure 3B).

**Figure 3:**
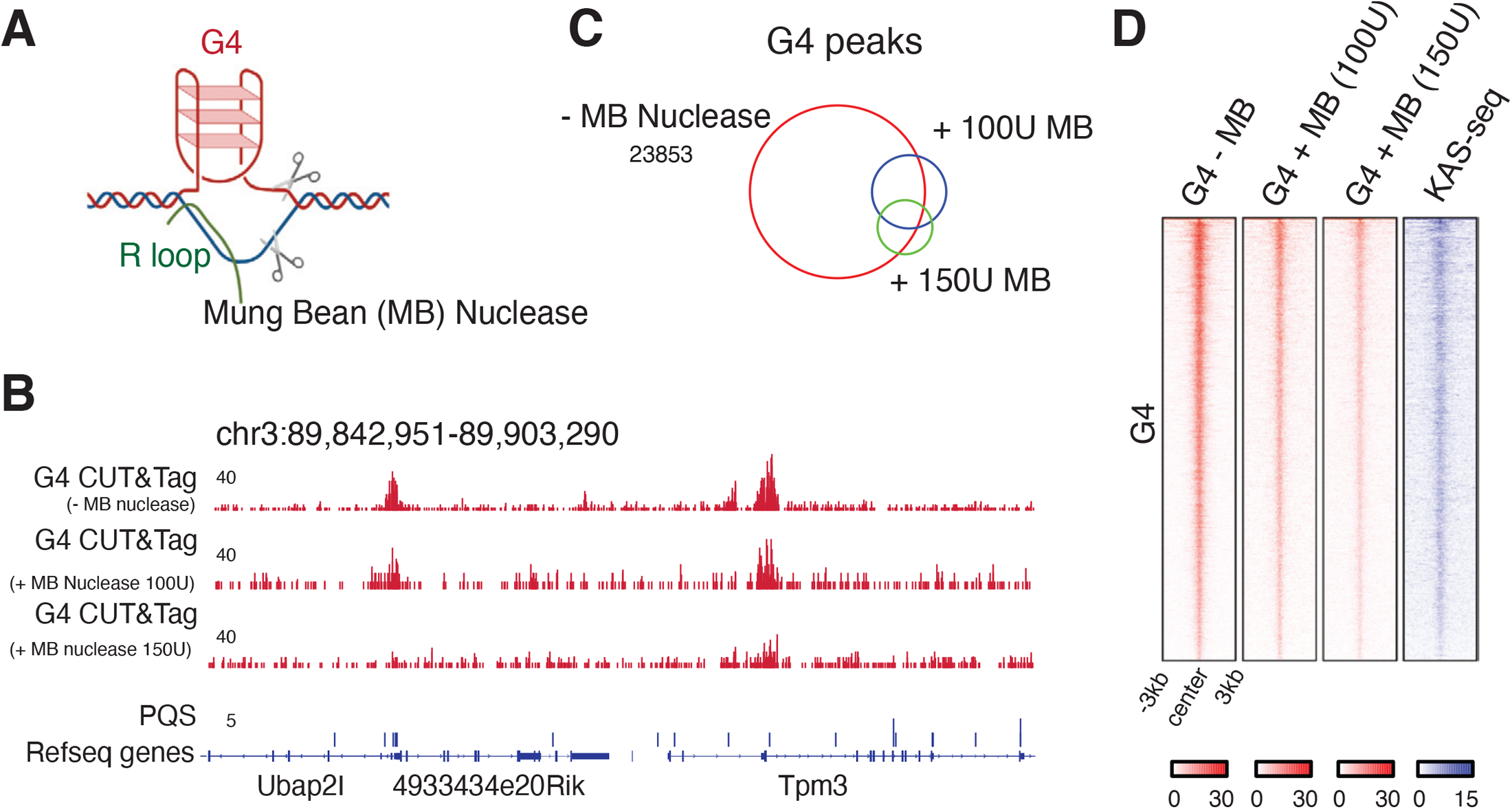
Single-strand specific endonuclease treatment. (**A**) Mung Bean (MB) nuclease treatment inhibits CUT&Tag amplification (**B**) Genome browser view showing G4 CUT&Tag signals in mES cells with or without Mung Bean nuclease pretreatment (**C**) Venn diagram showing overlap of G4 CUT&Tag peaks from native, 100U of MB treatment and 150U of MB treatment conditions. (**D**) Density heatmaps of native G4 CUT&Tag, 100U MB-treated G4 CUT&Tag,150U MB-treated G4 CUT&Tag, and single-stranded DNA from KAS-seq (50) at high-confidence G4 peak regions.

### G4s are paired with R loops genome-wide in mESC

DNA-RNA hybrids (R loops) formed by the opposing C-rich strand with nascent transcripts are thought to promote and stabilize G4 structures, and vice versa (16). Various R-loop mapping methods have been described to date, generating substantially different profiles: R-loops are most abundant in AT-rich regions in S1-DRIP-seq method (53), but most of the other DRIP-seq methods established a prevalence of R-loops at CG-rich regions (54-56). We thus wondered if R-loop can also be detected with the CUT&Tag protocol using an R-loop specific S9.6 monoclonal antibody (57). In line with a recent report validating the utility of S9.6 CUT&Tag for mapping R loops (58), we achieved a high signal-to-noise ratio and determined R loops genome-wide in mES cells. R-loops mapped with S9.6 CUT&Tag were highly prevalent at active promoters which also exhibited high RNA Pol II occupancy (Figure 4A) High-confidence R-loop CUT&Tag peaks were generated from the intersection of two replicates, and R-loop signals were found to be highly enriched at active promoters (34%) and enhancers (20%) (Figure 4B, Supplementary Figure 4A-D). Comparing G4 and R-loop signals by CUT&Tag in mES cells revealed that R-loop correlated well with G4 at TSS (Figure 4C), and 71% R-loop peaks overlapped with G4 peaks (Figure 4D). Plotting G4 and R-loop signal across shared, G4-only and R-loop only peaks revealed that the true overlap was greater (Figure 4C): most R-loop peaks showed some signal for G4 and vice versa. These results collectively demonstrate that G4 and R-loop show high co-occurrence and provide more evidence for the involvement of G4 in transcription regulation.

**Figure 4:**
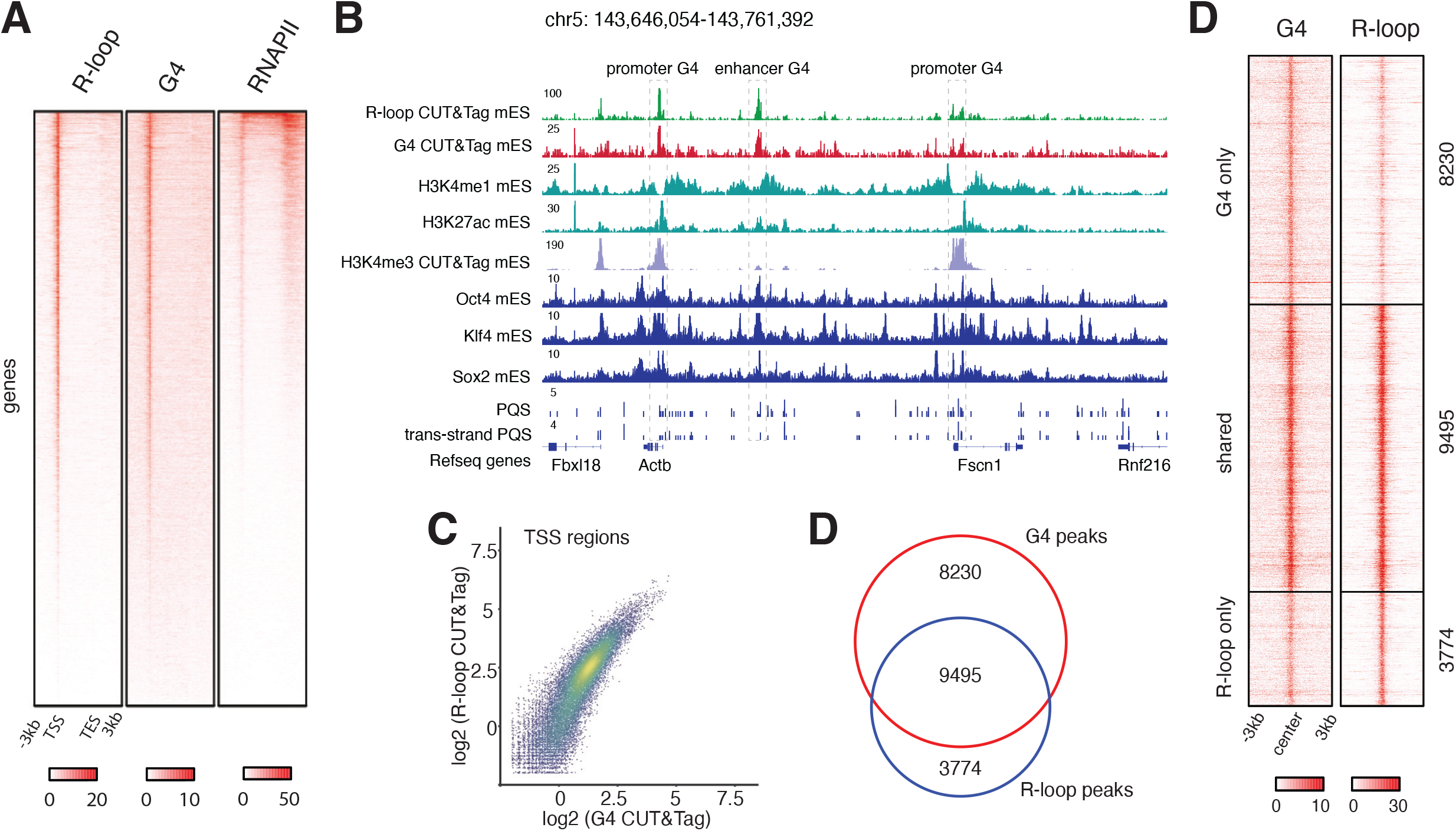
Genome-wide coincidence of R-loops and G4s. (**A**) Density heatmaps of R-loop CUT&Tag, G4 CUT&Tag, RNA Pol II ChIP-Seq (66) at genes. (**B**) Genome browser view showing coincidence of R-loop and G4 at promoter and enhancer regions. Scatterplot showing relationship of G4 CUT&Tag and R-loop CUT&Tag signal at TSS. Pearson correlation coefficient was calculated. (**D**) Intersection of G4 CUT&Tag and R-loop CUT&Tag peaks. (**D**) G4 and R-loop CUT&Tag density heatmaps for peaks classified as G4 only (n = 8230), both G4 and R-loop (n = 9495), or R-loop only (n = 3774).

## DISCUSSION

To understand G4 biology, genome-wide mapping methods are indispensable. Here, we demonstrate that CUT&Tag provides a reliable method for detecting G4s across human and mouse genomes. Compared to published methods, G4 CUT&Tag shows a higher fraction of peaks with canonical and non-canonical G4 motifs (Figure 1G, H), and validation with single-stranded nuclease pre-treatment demonstrates that the observed signal indeed arises from regions where the DNA duplex is melted (Figure 3A).

Limitations of the CUT&Tag procedure exist and are also relevant for mapping G4 structures: currently, the protocol crucially relies on native cells and has not been shown to work with fixed tissues, as has been demonstrated for G4 ChIP-Seq (36). While we demonstrate an excellent signal-to-noise ratio measured against other available methodologies, the background of a G4 mapping method should theoretically be zero for regions without any PQS. Our Mung Bean nuclease experiment shows that in the absence of G4 structures, the BG4 antibody still produces a background of random genomic tagmentation events (remaining reads shown in bottom track of Figure 3B). In contrast, omitting BG4 or anti-FLAG secondary antibodies in the CUT&Tag procedure lead to a strong reduction in global tagmentation efficiency, producing only low-complexity libraries (data not shown).

The lack of an input control for the CUT&Tag signal makes it difficult to interpret enrichment at repetitive sequences. G4s have been suggested to form at interspersed G-rich tandem repeat regions and telomeres in vivo, thus quantitation of G4 signals at such regions would be desirable as part of a genome-wide study. However we note that the proportion of repeat sequences in CUT&Tag data is variable between replicates and thus cannot be reliably quantified (data not shown).

Despite the abundance of G4-forming motifs present across human and mouse genomes, we find that formation of stable G4s requires open chromatin, as found at active promoters and enhancers. Interestingly, while canonical G4 motifs are more prevalent at promoters, we find that enhancers often do not feature the four GGG repeats on the same strand, and G4s may instead involve GGG repeats on both strands of the DNA duplex (Figure 2E). An alternative model, supported by *in silica* and *in vitra* data, suggests that G4s can fold in trans at chromatin loop anchors, e.g. with contribution of two GGG repeats by the promoter and two by the enhancer (59, 60). The existence of such trans-loop or ‘kissing’ loop G4s in cells is difficult to prove experimentally but it is intriguing to speculate that such G4s could stabilize promoter-enhancer interactions through a direct tethering of the DNA (59, 60). Interestingly, we find that only embryonic active enhancers feature non-canonical G4 motifs, whereas primed enhancers (carrying H3K4me1 but not H3K27ac) in mouse embryonic stem cells generally lack G4 motifs. This is an unexpected finding, suggesting that G4s could contribute uniquely to enhancer function in pluripotent embryonic stem cells. The underlying reason for such sequence-encoded bias is yet to be elucidated but we note that the pluripotency-associated transcription factor KLF4 recognizes a GpG dinucleotides within a G-rich context (61, 62). Thus, evolutionary selection for KLF4-binding motifs may have enriched embryonic enhancers in G4 motifs.

In summary, we have shown here that CUT&Tag provides a reliable and simple approach for genome-wide mapping of G4 structures and R loops, and we envision that the method will be generally useful for mapping of non-canonical DNA structures for which specific antibodies or specific binding modules are available.

## METHODS

### Cell culture

HEK293T and HaCaT cells were cultured in DMEM high glucose, GlutaMAX™ Supplement, pyruvate (LifeTechnologies, 10569010), 10% fetal bovine serum (Sigma, F7524) under standard conditions (10% CO2, 90% humidity, 37 °C). Mouse embryonic stem cells (mESCs) were cultured feeder-free in 0.1% gelatin-coated flasks (Sigma, G1890) under standard conditions (10% CO2, 90% humidity, 37 °C) in KnockOut DMEM (LifeTechnologies, 10829018), 2 mM Alanyl-glutamine (Sigma, G8541), 0.1 mM non-essential amino acids (Sigma, M7145), 15% fetal bovine serum (FBS) (Sigma, F7524), 0.1 mM -mercaptoethanol (Sigma, M3148), ESGRO Leukemia Inhibitory Factor (LIF) (Millipore, ESG1107), 1µM PD0325901 (PZ0162-25MG) and 3µM CHIR99021 (SML1046-25MG).

### Cleavage under targets and tagmentation

Recombinant BG4 antibody and Protein A-Tn5 (pA-Tn5) were purchased from the Protein Science Facility at the Department of Molecular Biochemistry and Biophysics at Karolinska Institutet. CUT&Tag experiment was performed as described previously (45) with minor modifications. Briefly, 1×10^5^ cells were harvested, washed with wash buffer (20mM HEPES pH 7.5, 150mM NaCl, 0.5mM spermidine), and immobilized to concanavalin A-coated beads (Bangs Laboratories, BP531) with incubation at room temperature for 10min. The beads-bound cells were incubated in 200µl of primary antibody buffer (wash buffer with 2mM EDTA and 0.05% digitonin) with 1:100 BG4 antibody or S9.6 (Millipore, MABE1095) antibody dilution at 4°C by rotating overnight. The next day, the primary antibody buffer was removed and cells were washed with 800µl of dig-wash buffer (wash buffer with 0.05% digitonin) three times. After washing, BG4 antibody-incubated cells were resuspended with 200ul of dig-wash buffer with 1:100 dilution of anti-FLAG antibody (Sigma, F1804) and incubated at room temperature for 1 hour by rotating. Cells were washed with 800µl of dig-wash buffer briefly three times to remove unbound antibodies. S9.6-treated or anti-FLAG treated cells were incubated with 1:100 dilution of anti-mouse antibody (Sigma, M7023) in 200µl of dig-wash buffer at room temperature for 1 hour by rotating.

After brief wash with dig-wash buffer as described, cells were resuspended in 200ul of dig-300 buffer (20mM HEPES pH 7.5, 300mM NaCl and 0.5mM spermidine, 0.01% digitonin) with 1:200 dilution of pA-Tn5 adapter complex and incubated at room temperature for 1 hour by rotating. pA-Tn5-bound cells were washed with 800ul of dig-300 buffer three times, followed by tagmentation in 200ul of tagmentation buffer (dig-300 buffer with 10mM MgCl_2_) at 37°C for 1 hour. After tagmentation, 15mM EDTA, 500µg/ml proteinase K and 0.1% SDS were added and further incubated at 63°C for another 1 hour to stop tagmentation and digest protein. Genomic DNA was extracted and purified with DNA Clean & Concentrator-5 (Zymo research, D4013). To generate G4 or R-loop libraries, purified genomic DNA were mixed with the universal i5 primer and barcoded i7 primer, and amplified with NEBNext Ultra II Q5 Master Mix (NEB, M0544). The library PCR products were cleaned up with Agencourt AMPure XP beads (Beckman Coulter, A63881) and sequenced with Illumina Nextseq 500 instrument.

### Mapping pipeline

G4 and R-loop CUT&Tag, as well as published G4 ChIP-Seq and G4P ChIP-Seq data were processed as follows; reads were aligned with bowtie2 (v.2.3.5.1) (68) using --fast-local and samtools (v1.10) (69). BAM files were deduplicated with picard (v2.23.4) MarkDuplicates. Blacklisted regions were removed from the BAM file with bedtools (v2.29.2) intersect (70) using ENCODE blacklist bed files for mm9 or hg19. Normalized (RPGC, 1x Genome Coverage) coverage tracks were generated using deepTools (v3.3.2) bamCoverage (71) using parameters -binSize 5 --normalizeUsing RPGC. Peaks were called with MACS2 (72). High confidence peaks sets were generated by intersecting the peaks called for 3 replicates (G4s) or 2 replicates (R-loops).

### Quality metrics

FriP was calculated with featureCounts (73). Fingerplots were generated with deepTools (v3.3.2) plotFingerprint (71).

### Bioinformatic analysis

Analyses were generated from normalized bigwig files. Profile plots and heatmaps were generated with SeqPlots (74). Scatterplots were generated using wigglescout (https://github.com/cnluzon/wigglescout/), an R library for bigWig genomics data visualization.

### G-quadruplex motif search

Genomic sequences of mm9 and hg19 were subjected to two classes of G4 pattern matching, interstrand motifs (75) and intrastrand motifs (19). For interstrand motifs, the canonical G4 motif G_3+_L_1-7_ was expanded to the opposite strand in 8 combinations. Let A = G_3+_ and B = C_3+_, the 8 patterns are AAAA, AAAB, AABA, AABB, ABAA, ABAB, ABBA and ABBB. The canonical intrastrand G4 pattern was the same as AAAA; the extra intrastrand patterns were extended canonical PQS (Putative G-Quadruplex Sequences) G_3+_L_1-12_, and two-tetrads PQS G_2_L_1-12_. Regular expression was applied, for example two-tetrads PQS was ‘[gG]{2}\\W{1,12}{3,}[gG]{2}’. Motif genome coverage was generated with R package “rtracklayer 1.46.0” either by the predicted binary hits or stack of occurrences, and exported as bigwig.

## Data availability

G4 ChIP-Seq, G4P ChIP-Seq and mES public ChIP-seq were downloaded from GEO: GSM2035780 (HaCaT G4 ChIP-seq), GSM2035782 (HaCaT G4 ChIP-seq, input), GSM3907020 (HEK293T G4P-ChIP), GSM3907021 (HEK293T G4P-ChIP, input), GSM1917298 (mES Ring1b ChIP-seq), SRR10349547 (mES KAS-seq), GSM1173376 (mES S2 PolII ChIP-seq), GSM4205678 (mES H3K27ac ChIP-seq), GSM4303796 (mES H3K4me1 ChIP-seq), GSM4661960 (mES ATAC-seq), GSM1127953 (mES Bisulfite-seq), GSM2582392 (mES H3K9me3 ChIP-seq). CUT&Tag data generated for this manuscript has been deposited at the Gene Expression Omnibus under GSE173103.

## Code availability

Associated code is available at https://github.com/elsasserlab/G4

## ACKNOWLEDGEMENTS

Bioinformatics analyses were performed on resources provided by the Swedish National Infrastructure for Computing (SNIC) at Uppmax server (projects SNIC 2020/15-9, SNIC 2020/6-3, uppstore2018208, SNIC 2018/3-669, sllstore2017057, SNIC 2017/1-508). We thank the Protein Science Facility at the Department of Molecular Biochemistry and Biophysics at Karolinska Institutet for producing pA-Tn5 and BG4 antibody proteins. We thank members of the Elsässer lab for comments and help with experiments and analysis.

S.J.E acknowledges funding by the Karolinska Institutet SFO for Molecular Biosciences, Vetenskapsrådet Junior Researcher Grant (2015-04815), H2020 ERC Starting Grant (715024 RAPID), Åke Wibergs Stiftelse (M15-0275), Cancerfonden (2015/430). J.L. and R.S. acknowledge funding from the Chinese Scholarship Council.

## AUTHOR CONTRIBUTIONS

J.L. and S.J.E. conceived study. J.L. performed all experiments. R.S. helped analyzing data. J.L. and S.J.E. analyzed data, generated figures and wrote the manuscript.

---

**Supplementary Figure 1 -.**
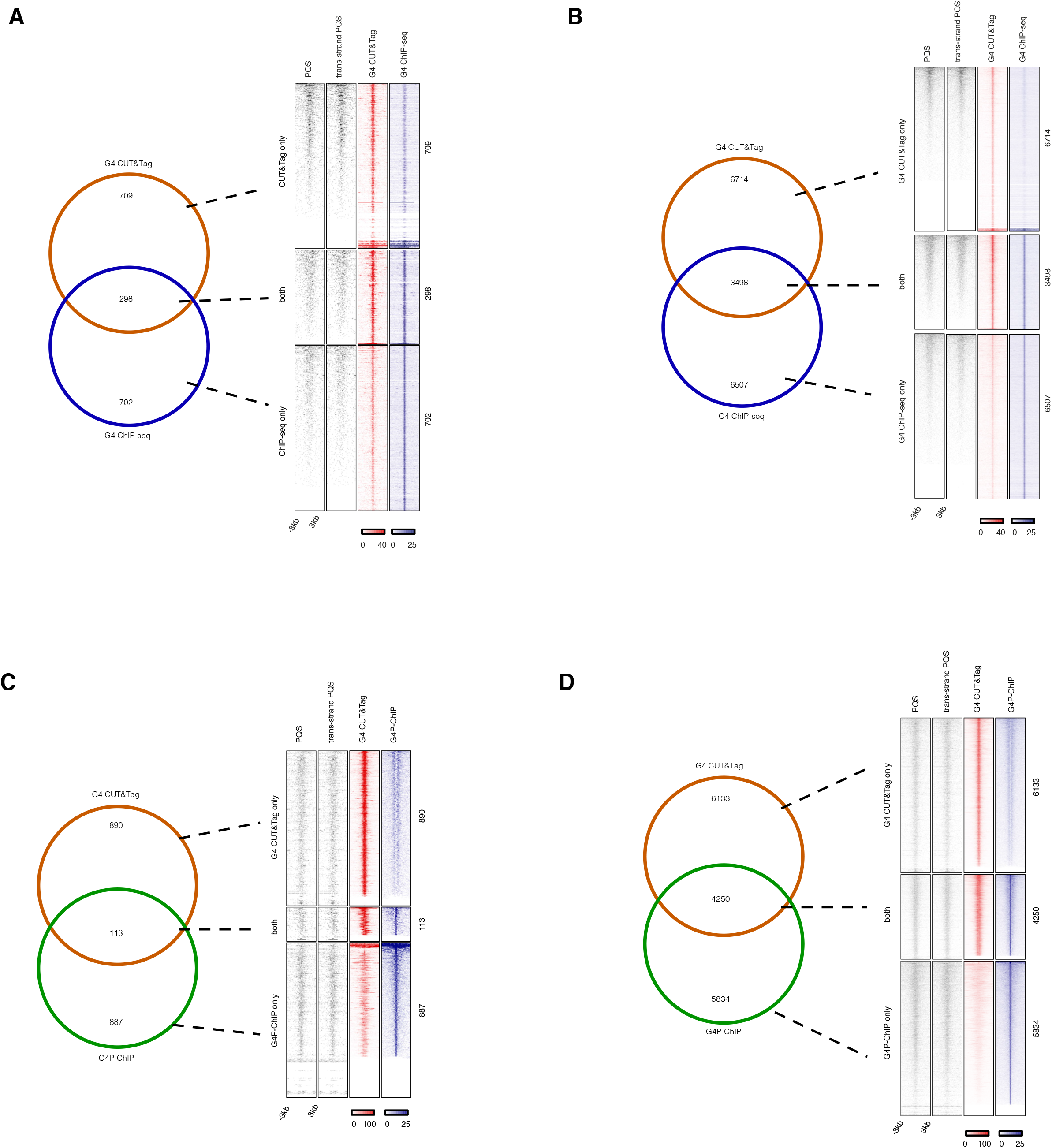
Comparison of G4 CUT&Tag to other G4 mapping methods, G4 ChIP-seq and G4P-ChIP. (**A**) Overlap of top 1000 G4 CUT&Tag and G4 ChIP-seq peaks. Total PQS, trans-strand PQS, G4 CUT&Tag and G4 ChIP-seq density heatmaps for peaks classified as G4 CUT&Tag only (n = 709), both G4 CUT&Tag and G4 ChIP-seq (n = 298) or G4 ChIP-seq only (n = 702). (**B**) Overlap of top 10,000 G4 CUT&Tag and G4 ChIP-seq peaks. Total PQS, trans-strand PQS, G4 CUT&Tag and G4 ChIP-seq density heatmaps for peaks classified as G4 CUT&Tag only (n = 6714), both G4 CUT&Tag and G4 ChIP-seq (n = 3498) or G4 ChIP-seq only (n = 6507). (**C**) Overlap of top 1000 G4 CUT&Tag and G4P-ChIP peaks. Total PQS, trans-strand PQS, G4 CUT&Tag and G4 ChIP-seq density heatmaps for peaks classified as G4 CUT&Tag only (n = 890), both G4 CUT&Tag and G4 ChIP-seq (n = 113) or G4 ChIP-seq only (n = 887). Overlap of top 10,000 G4 CUT&Tag and G4P-ChIP peaks. Total PQS, trans-strand PQS, G4 CUT&Tag and G4 ChIP-seq density heatmaps for peaks classified as G4 CUT&Tag only (n = 6133), both G4 CUT&Tag and G4 ChIP-seq (n = 4250) or G4 ChIP-seq only (n = 5834).

**Supplementary Figure 2:**
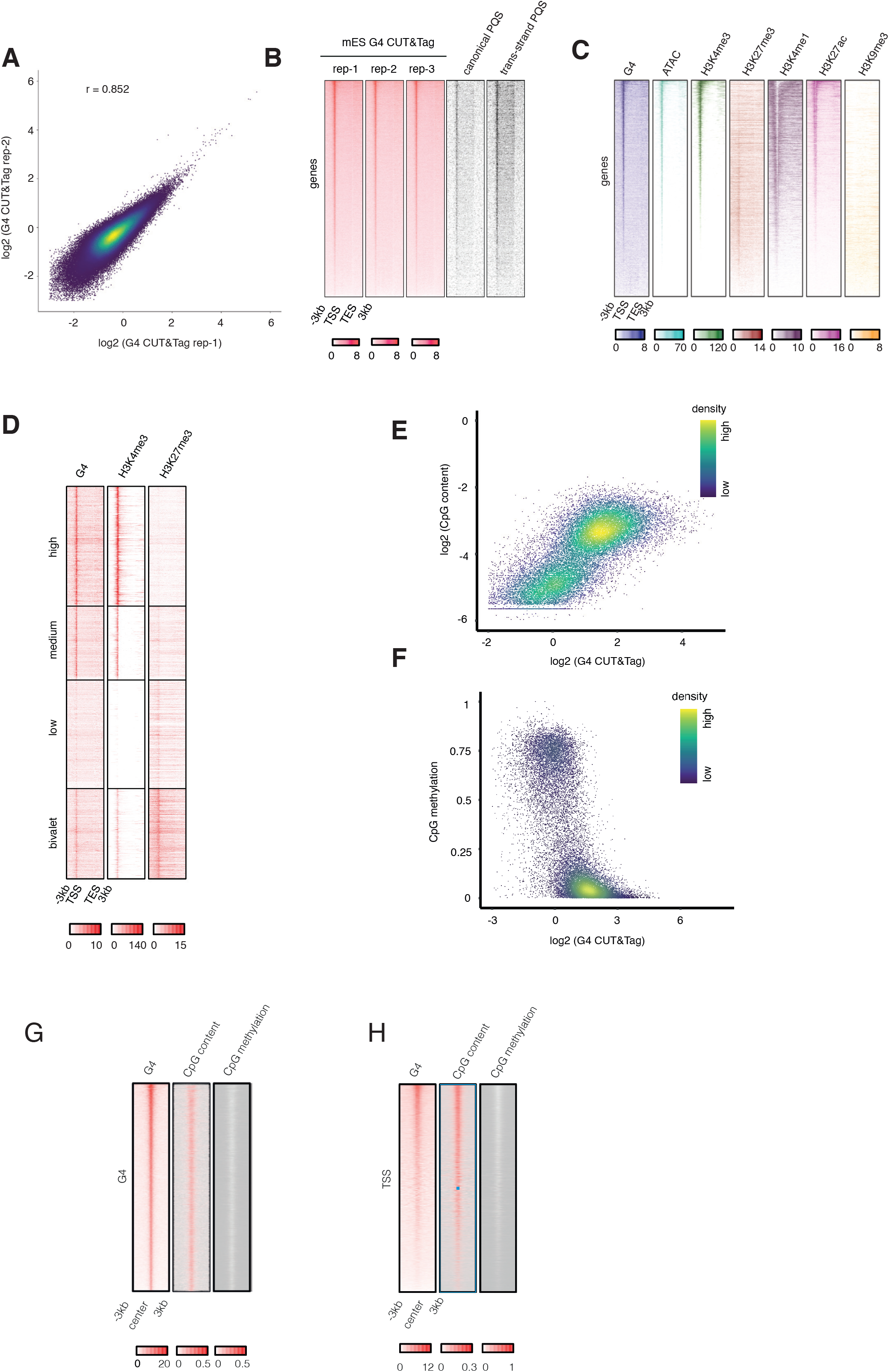
G4 landscape in mouse embryonic stem cells. (**A**) Scatterplot of 10kb windows genome-wide showing the reproducibility of G4 CUT&Tag signal between two mES replicates. Pearson correlation was calculated. (**B**) Density heatmaps showing the reproducibility of G4 CUT&Tag at gene-coding regions, and overlap with canonical and non-canonical (trans-strand) PQS. (**C**) Density heatmaps of G4 CUT&Tag,H3K4me3 and H3K27me3 CUT&Tag from the same cells, as well as published ATAC-seq (63), H3K4me1, H3K27ac, H3K9me3 ChIP-Seq (64). (**D**) Density heatmaps of G4, H3K4me3, H3K27me3 CUT&Tag at representative groups (∼2000 genes per group) of highly-expressed, mediumly-expressed, low/non-expressed and bivalent genes. (**E**) Scatterplot showing the correlation of G4 CUT&Tag signal and CpG content at TSS regions. (**F**) Scatterplot showing the anti-correlation of G4 CUT&Tag and CpG methylation (67) at TSS regions. (**G**) Density heatmaps of G4 CUT&Tag, CpG content and CpG methylation at G4 peaks. (**H**) Density heatmaps of G4 CUT&Tag, CpG content and CpG methylation at TSS regions.

**Supplementary Figure 3:**
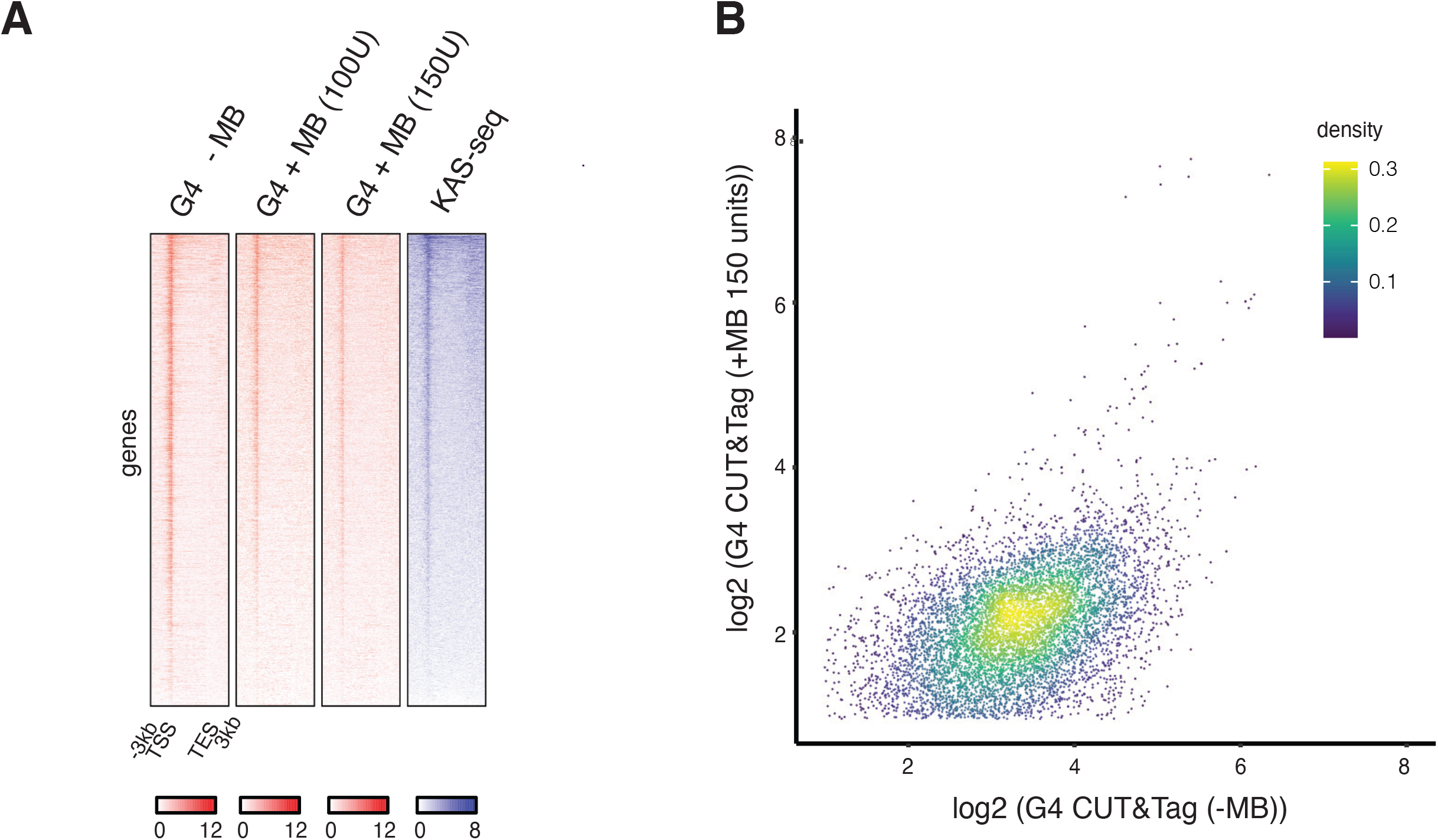
Single-strand specific endonuclease treatment. (**A**) Density heatmaps of native G4 CUT&Tag, 100U MB-treated G4 CUT&Tag,150U MB-treated G4 CUT&Tag and KAS-seq (50) at gene-coding regions. (**B**) Scatterplot of G4 peaks showing the relationship of native G4 CUT&Tag and 150U MB-treated G4 CUT&Tag.

**Supplementary figure 4:**
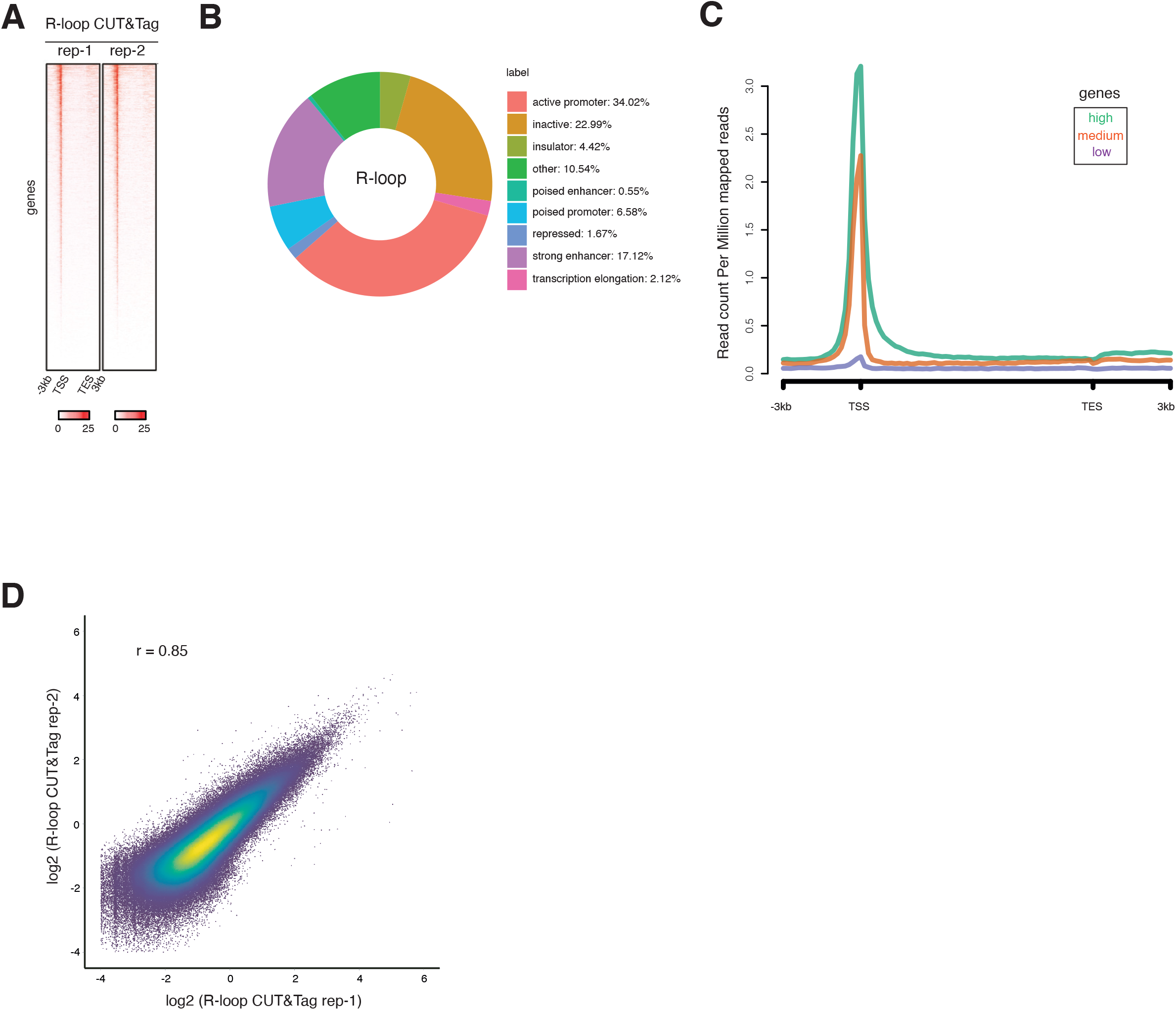
Genome-wide coincidence of R-loops and G4s. (**A**) Density heatmaps showing the reproducibility of R-loop CUT&Tag replicates at gene-coding regions. (**B**) Annotation of high-confidence R-loop peaks with different functional genomic features as defined by ChromHMM (65). (**C**) Average R-loop CUT&Tag signals at high-expressed, medium-expressed and low/non-expressed gene-coding regions. Gene body and 3kb upstream of TSS and 3kb downstream of TES is shown. (**D**) Scatterplot showing the reproducibility of R-loop CUT&Tag replicates. Pearson correlation was calculated.

## REFERENCES

1. Burge, S., Parkinson, G.N., Hazel, P., Todd, A.K. and Neidle, S. (2006) Quadruplex DNA: sequence, topology and structure. Nucleic Acids Res., 34, 5402–5415.

2. Rawal, P., Kummarasetti, V.B.R., Ravindran, J., Kumar, N., Halder, K., Sharma, R., Mukerji, M., Das, S.K. and Chowdhury, S. (2006) Genome-wide prediction of G4 DNA as regulatory motifs: role in Escherichia coli global regulation. Gename Res., 16, 644–655.

3. Schaffitzel, C., Berger, I., Postberg, J., Hanes, J., Lipps, H.J. and Pluckthun, A. (2001) In vitro generated antibodies specific for telomeric guanine-quadruplex DNA react with Stylonychia lemnae macronuclei. Prac Natl Acad Sci USA, 98, 8572–8577.

4. Paeschke, K., Simonsson, T., Postberg, J., Rhodes, D. and Lipps, H.J. (2005) Telomere end-binding proteins control the formation of G-quadruplex DNA structures in vivo. Nat. Struct. Mal. Bial., 12, 847–854.

5. Eddy, J. and Maizels, N. (2008) Conserved elements with potential to form polymorphic G-quadruplex structures in the first intron of human genes. Nucleic Acids Res., 36, 1321–1333.

6. Maizels, N. and Gray, L.T. (2013) The G4 genome. PLaS Genet., 9, e1003468.

7. Todd, A.K., Johnston, M. and Neidle, S. (2005) Highly prevalent putative quadruplex sequence motifs in human DNA. Nucleic Acids Res., 33, 2901–2907.

8. Huppert, J.L. and Balasubramanian, S. (2005) Prevalence of quadruplexes in the human genome. Nucleic Acids Res., 33, 2908–2916.

9. Fay, M.M., Lyons, S.M. and Ivanov, P. (2017) RNA G-Quadruplexes in Biology: Principles and Molecular Mechanisms. J. Mal. Bial., 429, 2127–2147.

10. Yang, S.Y., Lejault, P., Chevrier, S., Boidot, R., Robertson, A.G., Wong, J.M.Y. and Monchaud, D. (2018) Transcriptome-wide identification of transient RNA G-quadruplexes in human cells. Nat. Cammun., 9, 4730.

11. Siddiqui, A. Direct evidence for a G-quadruplex in a promoter region and its targeting with a small molecule to repress c-MYC transcription. Jain.

12. Cogoi, S. and Xodo, L.E. (2006) G-quadruplex formation within the promoter of the KRAS proto-oncogene and its effect on transcription. Nucleic Acids Res., 34, 2536–2549.

13. Rankin, S., Reszka, A.P., Huppert, J., Zloh, M., Parkinson, G.N., Todd, A.K., Ladame, S., Balasubramanian, S. and Neidle, S. (2005) Putative DNA quadruplex formation within the human c-kit oncogene. J. Am. Chem. Sac., 127, 10584–10589.

14. Wang, Y., Yang, J., Wild, A.T., Wu, W.H., Shah, R., Danussi, C., Riggins, G.J., Kannan, K., Sulman, E.P., Chan, T.A., et al. (2019) G-quadruplex DNA drives genomic instability and represents a targetable molecular abnormality in ATRX-deficient malignant glioma. Nat. Cammun., 10, 943.

15. Miglietta, G., Russo, M. and Capranico, G. (2020) G-quadruplex-R-loop interactions and the mechanism of anticancer G-quadruplex binders. Nucleic Acids Res., 48, 11942–11957.

16. De Magis, A., Manzo, S.G., Russo, M., Marinello, J., Morigi, R., Sordet, O. and Capranico, G. (2019) DNA damage and genome instability by G-quadruplex ligands are mediated by R loops in human cancer cells. Prac Natl Acad Sci USA, 116, 816–825.

17. Read, M.A. and Neidle, S. (2000) Structural characterization of a guanine-quadruplex ligand complex. Biachemistry, 39, 13422–13432.

18. Harikrishna, S., Kotaru, S. and Pradeepkumar, P.I. (2017) Ligand-induced conformational preorganization of loops of c-MYC G-quadruplex DNA and its implications in structure-specific drug design. Mal. Biasyst., 13, 1458–1468.

19. Marsico, G., Chambers, V.S., Sahakyan, A.B., McCauley, P., Boutell, J.M., Antonio, M.D. and Balasubramanian, S. (2019) Whole genome experimental maps of DNA G-quadruplexes in multiple species. Nucleic Acids Res., 47, 3862–3874.

20. Di Antonio, M., Ponjavic, A., Radzevicius, A., Ranasinghe, R.T., Catalano, M., Zhang, X., Shen, J., Needham, L.-M., Lee, S.F., Klenerman, D., et al. (2020) Single-molecule visualization of DNA G-quadruplex formation in live cells. Nat. Chem., 12, 832–837.

21. Grand, C.L., Han, H., Munoz, R.M. and Weitman., S. (2002) The Cationic Porphyrin TMPyP4 Down-Regulates c-MYC and Human Telomerase Reverse Transcriptase Expression and Inhibits Tumor Growth in Vivo 1 This .. Malecular cancer ..

22. Mikami, Y. Antitumor activity of G-quadruplex-interactive agent TMPyP4 in K562 leukemic cells. Teraa.

23. Muller, S., Sanders, D.A., Di Antonio, M., Matsis, S., Riou, J.-F., Rodriguez, R. and Balasubramanian, S. (2012) Pyridostatin analogues promote telomere dysfunction and long-term growth inhibition in human cancer cells. Org. Biamal. Chem., 10, 6537–6546.

24. Rodriguez, R., Miller, K.M., Forment, J.V., Bradshaw, C.R., Nikan, M., Britton, S., Oelschlaegel, T., Xhemalce, B., Balasubramanian, S. and Jackson, S.P. (2012) Small-molecule-induced DNA damage identifies alternative DNA structures in human genes. Nat. Chem. Bial., 8, 301–310.

25. Guilbaud, G., Murat, P., Recolin, B., Campbell, B.C., Maiter, A., Sale, J.E. and Balasubramanian, S. (2017) Local epigenetic reprogramming induced by G-quadruplex ligands. Nat. Chem., 9, 1110–1117.

26. Drygin, D., Siddiqui-Jain, A., O’Brien, S., Schwaebe, M., Lin, A., Bliesath, J., Ho, C.B., Proffitt, C., Trent, K., Whitten, J.P., et al. (2009) Anticancer activity of CX-3543: a direct inhibitor of rRNA biogenesis. Cancer Res., 69, 7653–7661.

27. Papadopoulos, K.P. and Northfelt., D.W. (2007) Phase I clinical trial of CX-3543, a protein-rDNA quadruplex inhibitor. Jaurnal af Clinical ..

28. Drygin, D., Whitten, J., Rice, W., O’Brien, S. and Schwaebe., M. (2008) Quarfloxin (CX-3543) disrupts the Nucleolin/rDNA quadruplex complexes, inhibits the elongation by RNA Polymerase I and exhibits potent antitumor activity in models ..

29. Rice, W.G., Lim, J.K.C., Schwaebe, M.K. and Siddiqui, A. Design of CX-3543, a novel multi-targeting antitumor agent. Jain..

30. Xu, H., Di Antonio, M., McKinney, S., Mathew, V., Ho, B., O’Neil, N.J., Santos, N.D., Silvester, J., Wei, V., Garcia, J., et al. (2017) CX-5461 is a DNA G-quadruplex stabilizer with selective lethality in BRCA1/2 deficient tumours. Nat. Cammun., 8, 14432.

31. Drygin, D., Lin, A., Bliesath, J., Ho, C.B., O’Brien, S.E., Proffitt, C., Omori, M., Haddach, M., Schwaebe, M.K., Siddiqui-Jain, A., et al. (2011) Targeting RNA polymerase I with an oral small molecule CX-5461 inhibits ribosomal RNA synthesis and solid tumor growth. Cancer Res., 71, 1418–1430.

32. Haddach, M., Schwaebe, M.K., Michaux, J., Nagasawa, J., O’Brien, S.E., Whitten, J.P., Pierre, F., Kerdoncuff, P., Darjania, L., Stansfield, R., et al. (2012) Discovery of CX-5461, the First Direct and Selective Inhibitor of RNA Polymerase I, for Cancer Therapeutics. ACS Med. Chem. Lett., 3, 602–606.

33. Colis, L., Peltonen, K., Sirajuddin, P., Liu, H., Sanders, S., Ernst, G., Barrow, J.C. and Laiho, M. (2014) DNA intercalator BMH-21 inhibits RNA polymerase I independent of DNA damage response. Oncatarget, 5, 4361–4369.

34. Peltonen, K., Colis, L., Liu, H., Jaamaa, S., Zhang, Z., Af Hallstrom, T., Moore, H.M., Sirajuddin, P. and Laiho, M. (2014) Small molecule BMH-compounds that inhibit RNA polymerase I and cause nucleolar stress. Mal. Cancer Ther., 13, 2537–2546.

35. Fu, X., Xu, L., Qi, L., Tian, H., Yi, D., Yu, Y., Liu, S., Li, S., Xu, Y. and Wang, C. (2017) BMH-21 inhibits viability and induces apoptosis by p53-dependent nucleolar stress responses in SKOV3 ovarian cancer cells. Oncal. Rep., 38, 859–865.

36. Hansel-Hertsch, R., Simeone, A., Shea, A., Hui, W.W.I., Zyner, K.G., Marsico, G., Rueda, O.M., Bruna, A., Martin, A., Zhang, X., et al. (2020) Landscape of G-quadruplex DNA structural regions in breast cancer. Nat. Genet., 52, 878–883.

37. Papadopoulou, C., Guilbaud, G., Schiavone, D. and Sale, J.E. (2015) Nucleotide Pool Depletion Induces G-Quadruplex-Dependent Perturbation of Gene Expression. Cell Rep., 13, 2491–2503.

38. Lerner, L.K. and Sale, J.E. (2019) Replication of G quadruplex DNA. Genes (Basel), 10.

39. Hansel-Hertsch, R., Spiegel, J., Marsico, G., Tannahill, D. and Balasubramanian, S. (2018) Genome-wide mapping of endogenous G-quadruplex DNA structures by chromatin immunoprecipitation and high-throughput sequencing. Nat. Pratac., 13, 551–564.

40. Mao, S.-Q., Ghanbarian, A.T., Spiegel, J., Martinez Cuesta, S., Beraldi, D., Di Antonio, M., Marsico, G., Hansel-Hertsch, R., Tannahill, D. and Balasubramanian, S. (2018) DNA G-quadruplex structures mold the DNA methylome. Nat. Struct. Mal. Bial., 25, 951–957.

41. Hansel-Hertsch, R., Beraldi, D., Lensing, S.V., Marsico, G., Zyner, K., Parry, A., Di Antonio, M., Pike, J., Kimura, H., Narita, M., et al. (2016) G-quadruplex structures mark human regulatory chromatin. Nat. Genet., 48, 1267–1272.

42. Zheng, K.-W., Zhang, J.-Y., He, Y., Gong, J.-Y., Wen, C.-J., Chen, J.-N., Hao, Y.-H., Zhao, Y. and Tan, Z. (2020) Detection of genomic G-quadruplexes in living cells using a small artificial protein. Nucleic Acids Res., 48, 11706–11720.

43. Biffi, G., Tannahill, D., McCafferty, J. and Balasubramanian, S. (2013) Quantitative visualization of DNA G-quadruplex structures in human cells. Nat. Chem., 5, 182–186.

44. Liu, H.-Y., Zhao, Q., Zhang, T.-P., Wu, Y., Xiong, Y.-X., Wang, S.-K., Ge, Y.-L., He, J.-H., Lv, P., Ou, T.-M., et al. (2016) Conformation Selective Antibody Enables Genome Profiling and Leads to Discovery of Parallel G-Quadruplex in Human Telomeres. Cell Chem. Bial., 23, 1261–1270.

45. Kaya-Okur, H.S., Wu, S.J., Codomo, C.A., Pledger, E.S., Bryson, T.D., Henikoff, J.G., Ahmad, K. and Henikoff, S. (2019) CUT&Tag for efficient epigenomic profiling of small samples and single cells. Nat. Cammun., 10, 1930.

46. Schlesinger, S. and Meshorer, E. (2019) Open chromatin, epigenetic plasticity, and nuclear organization in pluripotency. Dev. Cell, 48, 135–150.

47. Armas, P. and Calcaterra, N.B. (2018) G-quadruplex in animal development: Contribution to gene expression and genomic heterogeneity. Mech. Dev., 154, 64–72.

48. Viktorovskaya, O., Chuang, J., Jain, D., Reim, N.I., Lpez-Rivera, F., Murawska, M., Spatt, D., Churchman, L.S., Park, P.J. and Winston, F. (2021) Essential histone chaperones collaborate to regulate transcription and chromatin integrity. Genes Dev., 10.1101/gad.348431.121.

49. Cruz-Molina, S., Respuela, P., Tebartz, C., Kolovos, P., Nikolic, M., Fueyo, R., van Ijcken, W.F.J., Grosveld, F., Frommolt, P., Bazzi, H., et al. (2017) PRC2 Facilitates the Regulatory Topology Required for Poised Enhancer Function during Pluripotent Stem Cell Differentiation. Cell Stem Cell, 20, 689-705.e9.

50. Wu, T., Lyu, R., You, Q. and He, C. (2020) Kethoxal-assisted single-stranded DNA sequencing captures global transcription dynamics and enhancer activity in situ. Nat. Methads, 17, 515–523.

51. Kouzine, F., Wojtowicz, D., Baranello, L., Yamane, A., Nelson, S., Resch, W., Kieffer-Kwon, K.-R., Benham, C.J., Casellas, R., Przytycka, T.M., et al. (2017) Permanganate/S1 Nuclease Footprinting Reveals Non-B DNA Structures with Regulatory Potential across a Mammalian Genome. Cell Syst., 4, 344-356.e7.

52. Kowalski, D., Kroeker, W.D. and Laskowski, M. (1976) Mung bean nuclease I. Physical, chemical, and catalytic properties. Biachemistry, 15, 4457–4463.

53. Wahba, L., Costantino, L., Tan, F.J., Zimmer, A. and Koshland, D. (2016) S1-DRIP-seq identifies high expression and polyA tracts as major contributors to R-loop formation. Genes Dev., 30, 1327–1338.

54. Ginno, P.A., Lott, P.L., Christensen, H.C., Korf, I. and Chedin, F. (2012) R-loop formation is a distinctive characteristic of unmethylated human CpG island promoters. Mal. Cell, 45, 814–825.

55. Ginno, P.A., Lim, Y.W., Lott, P.L., Korf, I. and Chedin, F. (2013) GC skew at the 5’ and 3’ ends of human genes links R-loop formation to epigenetic regulation and transcription termination. Gename Res., 23, 1590–1600.

56. Chen, L., Chen, J.-Y., Zhang, X., Gu, Y., Xiao, R., Shao, C., Tang, P., Qian, H., Luo, D., Li, H., et al. (2017) R-ChIP Using Inactive RNase H Reveals Dynamic Coupling of R-loops with Transcriptional Pausing at Gene Promoters. Mal. Cell, 68, 745-757.e5.

57. Boguslawski, S.J., Smith, D.E., Michalak, M.A., Mickelson, K.E., Yehle, C.O., Patterson, W.L. and Carrico, R.J. (1986) Characterization of monoclonal antibody to DNA.RNA and its application to immunodetection of hybrids. J. Immunal. Methads, 89, 123–130.

58. Wang, K., Wang, H., Li, C., Yin, Z., Xiao, R., Li, Q., Xiang, Y., Wang, W., Huang, J., Chen, L., et al. (2021) Genomic profiling of native R loops with a DNA-RNA hybrid recognition sensor. Sci. Adv., 7.

59. Hegyi, H. (2015) Enhancer-promoter interaction facilitated by transiently forming G-quadruplexes. Sci. Rep., 5, 9165.

60. Williams, J.D., Houserova, D., Johnson, B.R., Dyniewski, B., Berroyer, A., French, H., Barchie, A.A., Bilbrey, D.D., Demeis, J.D., Ghee, K.R., et al. (2020) Characterization of long G4-rich enhancer-associated genomic regions engaging in a novel loop:loop “G4 Kissing” interaction. Nucleic Acids Res., 48, 5907–5925.

61. Jia, C., Carson, M.B., Wang, Y., Lin, Y. and Lu, H. (2014) A new exhaustive method and strategy for finding motifs in ChIP-enriched regions. PLaS ONE, 9, e86044.

62. Sun, W., Hu, X., Lim, M.H.K., Ng, C.K.L., Choo, S.H., Castro, D.S., Drechsel, D., Guillemot, F., Kolatkar, P.R., Jauch, R., et al. (2013) TherMos: Estimating protein-DNA binding energies from in vivo binding profiles. Nucleic Acids Res., 41, 5555–5568.

63. Navarro, C., Lyu, J., Katsori, A.-M., Caridha, R. and Elsasser, S.J. (2020) An embryonic stem cell-specific heterochromatin state promotes core histone exchange in the absence of DNA accessibility. Nat. Cammun., 11, 5095.

64. Zhang, T., Zhang, Z., Dong, Q., Xiong, J. and Zhu, B. (2020) Histone H3K27 acetylation is dispensable for enhancer activity in mouse embryonic stem cells. Gename Bial., 21, 45.

65. Ernst, J. and Kellis, M. (2012) ChromHMM: automating chromatin-state discovery and characterization. Nat. Methads, 9, 215–216.

66. Bunch, H., Zheng, X., Burkholder, A., Dillon, S.T., Motola, S., Birrane, G., Ebmeier, C.C., Levine, S., Fargo, D., Hu, G., et al. (2014) TRIM28 regulates RNA polymerase II promoter-proximal pausing and pause release. Nat. Struct. Mal. Bial., 21, 876–883.

67. Marks, H., Kalkan, T., Menafra, R., Denissov, S., Jones, K., Hofemeister, H., Nichols, J., Kranz, A., Stewart, A.F., Smith, A., et al. (2012) The transcriptional and epigenomic foundations of ground state pluripotency. Cell, 149, 590–604.

68. Langmead, B. and Salzberg, S.L. (2012) Fast gapped-read alignment with Bowtie 2. Nat. Methads, 9, 357–359.

69. Li, H., Handsaker, B., Wysoker, A., Fennell, T., Ruan, J., Homer, N., Marth, G., Abecasis, G., Durbin, R. and 1000 Genome Project Data Processing Subgroup (2009) The Sequence Alignment/Map format and SAMtools. Biainfarmatics, 25, 2078–2079.

70. Quinlan, A.R. (2014) BEDTools: The Swiss-Army Tool for Genome Feature Analysis. Curr. Pratac. Biainfarmatics, 47, 11.12.1-34.

71. Ramirez, F., Dundar, F., Diehl, S., Gruning, B.A. and Manke, T. (2014) deepTools: a flexible platform for exploring deep-sequencing data. Nucleic Acids Res., 42, W187–91.

72. Liu, T. (2014) Use model-based Analysis of ChIP-Seq (MACS) to analyze short reads generated by sequencing protein-DNA interactions in embryonic stem cells. Methads Mal. Bial., 1150, 81–95.

73. Liao, Y., Smyth, G.K. and Shi, W. (2014) featureCounts: an efficient general purpose program for assigning sequence reads to genomic features. Biainfarmatics, 30, 923–930.

74. Stempor, P. and Ahringer, J. (2016) SeqPlots - Interactive software for exploratory data analyses, pattern discovery and visualization in genomics. [version 1; peer review: 2 approved, 1 approved with reservations]. Wellcame Open Res., 1, 14.

75. Kudlicki, A.S. (2016) G-Quadruplexes Involving Both Strands of Genomic DNA Are Highly Abundant and Colocalize with Functional Sites in the Human Genome. PLaS ONE, 11, e0146174.

